# Scarless circular mRNA-based CAR-T cell therapy elicits superior anti-tumor efficacy

**DOI:** 10.1101/2024.08.05.606578

**Authors:** Qinchao Hu, Hui Zhao, Kaicheng Zhou, Xianxin Hua, Xuyao Zhang

**Affiliations:** Department of Biological Medicines and Shanghai Engineering Research Center of Immunotherapeutics, School of Pharmacy, Fudan University, Shanghai 201203, China; Byterna therapeutics Ltd., Shanghai 201203, China; Department of Cancer Biology, Abramson Family Cancer Research Institute, Perelman School of Medicine at the University of Pennsylvania, Philadelphia, PA, USA

## Abstract

Messenger RNA (mRNA)-based transient expression of CAR shows optimal safety profiles and provides promising opportunities to address existing challenges of viral vector-based CAR-T therapies and to meet emerging medical needs in noncancerous indications. However, linear mRNAs are intrinsically unstable and thus just achieve compromised efficacy. Here, we engineered a permuted intron exon (PIE) platform to synthesize scarless circular mRNA (cmRNA) for potent CAR expression and long-lasting efficacy. cmRNA significantly increased amount and duration of anti-CD19 CAR expression on human T cells. cmRNA-based anti-CD19 CAR-T cells elicit superior anti-tumor efficacy over linear mRNA counterparts, demonstrated by parallel lines of evidence including *in vitro* specific cell-killing, cytokine release, transcriptomics patterns, and *in vivo* tumor elimination and survival benefit. We found that cmRNA-based anti-CD19 CAR-T efficiently eliminated target cells *in vivo* and provide long-lasting antitumor efficacy. These results suggested that cmRNA could be a potent platform for unleashing full potential of mRNA technologies in cell therapies.

## Introduction

Engineering T cells with chimeric antigen receptor (CAR) has proven to be a landmark strategy for immunotherapy^1^. Since 2017, several CAR-T cell therapies have been approved by FDA and they are reshaping the clinical treatment landscape of hematological malignancies. As a universal platform for highly-specific elimination of targeted cells recognized by the CAR, the therapeutical potentials of CAR-T are far beyond oncology^1,2^. Recent attempts are expending application frontiers of CAR-T to noncancerous indications such as autoimmune diseases^3–5^, cardiac fibrosis^6^, and aging^7–9^.

Despite these remarkable advances, currently approved CAR-T cell therapies are continually raising major safety concerns, including serious cytokine-release syndrome (CRS), immune effector cell-associated neurotoxicity syndrome (ICANS), on-target/off-tumor toxicity, and insertion mutation-related secondary cancers^10–16^. Many of these side effects can be largely attributed to the viral vector-based CAR engineering technologies which inevitably lead to vector-related side effects and risks from genome integration and permanent CAR expression on T cells^17^. Therefore, huge efforts have being dedicated to develop alternatives including non-viral platforms^18^, precise CAR genome integration (e.g., CRISPR-Cas)^19^, and mRNA-based CAR expression^20^, to overcome aforementioned challenges.

Among these emerging technologies, mRNA-based CAR expression represents unique advantages including the transient protein expression, non-viral delivery platforms (e.g., lipid nanoparticle, LNP), minimal risk of transgene integration, and versatile drug modalities (*ex vivo* mRNA CAR-T and *in vivo* mRNA CAR)^21–24^. It is worth emphasizing that the risk-benefit balance will be the utmost consideration for CAR-T therapies in noncancerous indications^1,25^, which recapitulates the unique advantages of mRNA-based CAR-T to overcome existing challenges and to meet emerging medical need. Besides, because of the simplicity, low cost, and acceptable safety profile, engineering T cells with CAR-encoding mRNAs provides an alternative strategy for development of off-the shell, low-cost, and safe CAR-T cell therapy.

Continuous efforts have been directed to develop both *ex vivo* and *in vivo* mRNA-based CAR-T therapies over the last decade^26–32^. *Ex vivo* mRNA CAR-T therapies have been evaluated in early-stage clinical trials for malignant pleural mesotheliomas^30^, pancreatic cancer^31^, melanoma and breast cancer^32^, and these studies collectively demonstrated the good safety and feasibility of mRNA-based CAR-T cell therapy. Recently, the *in vivo* mRNA CAR therapy candidate MT-302 has entered clinical trial (ClinicalTrials.gov: NCT05969041), representing exciting frontiers of *in vivo* mRNA-CAR therapies. Although mRNA-based CAR-T therapies have been clinically tested ten years ago, their clinical translations are largely limited due to compromised efficacy. The transient expression nature of mRNAs is a double-edged sword in CAR-T applications. The bottleneck is that the linear mRNAs are intrinsically unstable *in vivo*, which inevitably leads to short duration of efficacy. Unlike the linear conformation of mRNA, circular mRNAs (cmRNAs) are covalently closed circular RNA molecules that are more stable than linear mRNAs^33,34^ and could be engineered to translate more proteins in mammalian cells^35–37^. Therefore, cmRNA-based CAR could conceptually enable higher and more durable CAR expression on the cell membrane of functional immune cells than linear mRNAs, holding the potential to generate more effective anti-tumor immunity.

Although cmRNA serves as a promising platform for mRNA CAR-T therapies, *in vitro* synthesizing cmRNA with high yield and no extra sequence insertion is still challenging^38^. For instance, enzymatic ligases (such as T4 RNA ligase) are widely used circularization methods but they are usually inefficient for long RNA chains (e.g., mRNAs) and usually suffer from extra sequence append during the T7-polymerase-based IVT process (which is prone to randomly append one or more nucleotides into the 3’ end of synthesized linear mRNA)^39^. Permuted intron-exon (PIE)-based ribozyme self-splicing strategies have advantages in increasing circularization yield and reducing the sequence inaccuracy problem because the ends of precursor linear mRNA will be spliced and removed from produced cmRNAs. However, classical PIE usually inserts long sequence segments (up to 186nt, also called scars) into produced cmRNAs, which are indispensable for triggering the circularization reaction^35^. A recent study optimized the classical PIE system to shorten the scars as shorter as 27nt while kept the comparable circularization yield^40^. Nevertheless, those introduced scars are derived from viral PIE exons and their unforeseeable immunogenicity raise nonnegligible risk concerns for developing cmRNA-based therapeutics^41–43^. Recent studies attempted to optimize PIE to synthesize scarless cmRNA but they still suffer from compromised circularization yield^44,45^.

Therefore, in this study, we engineered a high-yield and scarless PIE (Hi-Scarless-PIE) for synthesizing scarless cmRNA coding CAR construct and conducted proof-of-concept experiments to evaluate scarless cmRNA-based CAR-T cells in *in vitro* and *in vivo* models. Our findings demonstrated that scarless cmRNA-based CAR-T cell therapy elicits superior anti-tumor efficacy and exhibits clinically-relevant advantages over linear mRNA-based CAR-T cells, suggesting the scarless cmRNA-based CAR expression (cmCAR) could be a promising T cell engineering platform for mRNA-based CAR-T cell therapies.

## MATERIALS AND METHODS

### Cell culture and transfections

HEK-293T, Raji, and K562 cell lines were purchased from the Cell Bank, Chinese Academy of Sciences (CAS) and the NALM-6-luciferase (NALM-6-luc) reporter cell line was purchased from IMMOCELL company (IML-012). HEK-293T cells were cultured in DMEM medium (MeilunBio, MA0212) supplemented with 10% FBS (Cytiva, SH30396.03). Raji, K562, and NALM-6-luc cells were cultured in RPMI-1640 medium (MeilunBio, MA0215) supplemented with 10% FBS. Human primary PBMC (Peripheral blood mononuclear cell) were obtained from the Hycells company and maintained in RPMI-1640 with 10% FBS. Unless otherwise specified, all cells were maintained at 37°C in a humidified 5% CO_2_ atmosphere. For Gaussia luciferase (GLuc) and human erythropoietin (hEPO) expression, equimolar quantities of each RNAs (equivalent to 100 ng) were transfected to HEK-293T cells per well in 96-well plate using Lipofectmine 3000 (Thermo Fisher Scientific, L3000001) according to the manufacturer’s instructions. For EGFP expression, equimolar quantities of RNAs (equivalent to 500 ng) were transfected to HEK-293T cell per well in 24-well plate using Lipofectmine 3000.

### Protein expression measurement

Luminescence from Gaussia luciferase was detected using Pierce Gaussia Luciferase Glow Assay Kit (Thermo Fisher Scientific, 16161) on day 1, 2, 3, 4, 5, 6, and 7 after transfection. For luminescence detection at different time points, cell culture medium was fully removed and replaced with fresh medium every day. For EGFP fluorescence detection, images were taken 24 h after transfection using Olympus (CKX53). Cell culture supernatant was collected at 24 h after transfection and subjected to ELISA assays to measure the hEPO expression using Human Erythropoietin ELISA Kit (Abcam, ab274397).

### Vector construction and mRNA preparations

The protein coding sequence, IRES, 5’-UTR, 3’-UTR, and the fragments of the group I intron were synthesized by TSINGKE and cloned into pUC57 plasmid with a T7 promoter (Supplementary Table 1). The linearized plasmids were used as a template for *in vitro* transcription (IVT) using a T7 High Yield RNA Synthesis Kit (New England Biolabs, E2040). For the modified linear RNA, uridine was completed replacement with N1-methylpseudouridine (m1Ψ). Reactions were treated with DNaseI (New England Biolabs, M0303) after IVT. The synthesized RNA was column purified with Monarch RNA Cleanup Kit (New England Biolabs, M2040). The linear RNA was capped using the mRNA Cap 2’-O-Methyltransferase (New England Biolabs, M0366) and Vaccinia Capping System (New England Biolabs, M2080) and added the poly(A) tails using *E. coli* Poly(A) Polymerase (New England Biolabs, M0276) according to the manufacturer’s instructions.

### mRNA circularization and purification

The column purified cmRNA precursors were circularized in T4 RNA ligase buffer (New England Biolabs, B0216) with 2mM GTP at 55°C for 15 min. Then the products were treated with *E. coli* Poly(A) Polymerase (New England Biolabs, M0276) according to the manufacturer’s instructions and column purified. The purified RNA products were then treated with RNase R (Applied Biological Materials, E049) according to the manufacturer’s instructions and column purified to get the enriched cmRNA. For cmRNA splice junction sequencing, the RNaseR-treated cmRNA was reversely transcribed into cDNA using RevertAid First Strand cDNA Synthesis Kit (Thermo Fisher Scientific, K1622) with random primers. The PCR reaction was performed with primers crossing the splice junction site and the PCR products were sequenced by Sanger sequencing. cmRNAs were further analyzed using capillary electrophoresis on QIAxcel Connect machine using QIAxcel RNA High Sensitivity Cartridge Kit (QIAGEN, 929112).

### Lipid nanoparticle (LNP) formulation and characterization

The ionizable lipid DLin-MC3-DMA, along with DSPC, DMG-PEG 2000, and cholesterol, was sourced from AVT Pharmaceutical Technology Company. In the formulation of MC3-LNP, the lipid mixture, composed of 50 mol% DLin-MC3-DMA, 10 mol% DSPC, 38.5 mol% cholesterol, and 1.5 mol% DMG-PEG, was prepared in ethanol. The RNA was suspended in a 50 mM sodium citrate buffer solution adjusted to pH 4.0. The LNP-mRNA complex was fabricated through the emulsion of the lipid organic phase and the RNA aqueous phase at a microfluidic laminar flow rate of 12 mL/min, with an aqueous-to-organic phase flow rate ratio of 3:1, utilizing a Rapid Nanomedicine System (INanoTM L). Subsequently, the emulsion underwent a dialysis process, which was carried out three times, each for a duration of 2 hours at 4°C against a PBS buffer solution with a pH of 7.4, utilizing a Pur-A-LyzerTM Mega Dialysis Kit (MilliporeSigma, PURG60020). The resulting LNP formulation was concentrated using an Amicon Ultra 50 K MWCO centrifugal filter device (Merck Millipore, UFC905096) and subsequently passed through a 0.22-μm syringe filter (Millipore, SLGPR33RB) to ensure sterility and purity. Particle size distribution of the LNPs was ascertained via dynamic light scattering, as measured with a Zetasizer Nano ZS apparatus (Malvern Panalytical). The encapsulation efficiency of RNA was quantified using a Quant-it RiboGreen RNA Assay Kit (Thermo Fisher Scientific, R11490), following the protocol outlined by the manufacturer.

### Linear mRNA or cmRNA-based CAR T-cell production

Human PBMCs were cultured in RPMI-1640 supplemented with 10% FBS and activated with Dynabeads^TM^ Human T-Activator CD3/CD28 (Gibco, 11132D). One day post activation, IL-2 (PEPROTECH, 200-02) was supplemented to a final concentration of 300 U/mL and T cell density was maintained between 0.5 × 10^6^ - 1 × 10^6^ cells/mL. Beads were removed at day 3. Human T-cells pellets were collected and re-suspended in an electroporation buffer Opti-MEM (Gibco, 31985062), and the cell concentration was adjusted to be 8 × 10^7^ cells/ml. 250 µL of the cell suspension was mixed with equimolar quantities of linear mRNA and circular mRNA (equivalent to 20 µg) carefully prior to transfection. The cell-mRNA mixture was transferred into a 4 mm electroporation cuvette (BTX), and electroporation was performed using ECM2001+ system. Upon completion of electroporation, the cells were transferred to the pre-warmed culture medium for further cultivation.

### Flow cytometric analysis

Death cells were excluded using Zombie Violet^TM^ Fixable Viability Kit (BioLegend, 423114) following the manufacturer’s guidelines. CAR expression was determined by incubating CAR-T cells with PE-Labeled human CD19 (20-291) protein (ACRO, CD9-HP2H3) for 30 min at room temperature in FACS Buffer. Peripheral blood was obtained from retro-orbital bleeding and stained for the presence of total human T cells (anti-CD3), CAR-T cells (CD19 protein), memory T cells (anti-CD45RA (Biolegend, 304112), anti-CD62L (Biolegend, 304822)). Single-cell suspension was prepared for analysis by CytoFlex S flow cytometer (Beckman). More than 10000 events were acquired and analyzed in each sample.

### *In vitro* cytotoxicity

Cytotoxicity assays were conducted at the first day and fourth day following electroporation completion. CD19-positive (NALM-6 and Raji) and CD19-negative (K-562) cell lines were selected as target cells. Untransfected T cells, linear mRNA-CD19-CAR-T cells or cmRNA-CD19-CAR-T cells were incubated with target cells in flat-bottom 24-well plates for 16 h at an E:T of 1:1, 2:1 and 5:1. Then, the cells were collected and stained with Zombie Violet^TM^ Dye, anti-CD19 antibody and anti-CD3 antibody flow cytometry analysis. The lysis of tumor cells was determined by the summary (means ± SDs) of the following formula: Percentage of lysis (%) =100 × (1 - (CD19^+^ cell fraction of the indicated T cell group/CD19^+^ cell fraction in control T cell group)). Cytokines in the supernatants of cytotoxicity assay were analyzed using Human IFN-γ (Multi Sciences, EK180-96), TNF-ɑ (Multi Sciences, EK182-96) and Human IL-2 (Multi Sciences, EK102) ELISA kits according to the manufacturer’s instructions.

### Bulk RNA-seq experiments and bioinformatics analysis

Control T cells (untransfected), Linear mRNA CAR-T or cmRNA CAR-T cells (n=3) were harvested at day 2 and day 4 post electroporation transfections. Subsequently, the total RNA was extracted from cell samples using TRIzol Reagent (Thermo Fisher Scientific, 15596026CN) for preparing RNA libraries using the VVAHTS® Universal V8 RNA-seq Library Prep Kit for Illumina (Vazyme, NR605-0). Sequencing was performed using the Illumina NovaSeq 6000 platform.

To perform the preprocessing of raw data, the Trim galore (version 0.4.5) was used to remove aptamers and low-quality bases from raw RNAseq reads for each fastq file of each sample. Next, the trimmed reads were aligned to the reference genome (refdata-gex-GRCh38-2020-A) using STAR (Spliced Transcripts Alignment to a Reference, version 2.7.10b)^46^, taking into account splice alignment. Finally, the STAR-generated bam files were quantified using featureCounts (version 2.0.6)^47^ count the mapped reads for each gene, which generate a read count matrix for all samples over all genes. The downstream differential gene expression (DEG) analysis was performed based on the read count matrix using DESeq2 R package (v1.16.1)^48^. To calculate the TPM expression matrix, the rnanorm (version 1.5.1) was used based on the refdata-gex-GRCh38-2020-A reference genome file. The refdata-gex-GRCh38-2020-A reference file was obtained from https://www.10xgenomics.com/.

Differential expressed genes between CAR-T cells and control T cells were selected based on the following rules: the absolute value of log2 transformed Fold Change (log2FC) >= 1.0 and the FDR-adjusted P value (adjP) <= 0.05. DEG analysis was performed for multiple group comparisons: Linear-vs-UTD, Circular-vs-UTD, and Circular-vs-Linear at day 2 and day 4 experiments. When directly comparing cmCAR-T with mCAR-T, the threshold of log2FC was set as 0.5 to capture more significantly regulated genes (FDR-adjusted P value (adjP) <= 0.05). Gene Ontology (GO) and pathway enrichment analysis was conducted using Enrichr-KG webserver^49^ using GO_Biological_Process_2021, KEGG_2021_Human, and Reactome_2022 databases. T cell functional signature genes were collected from recent literature^50,51^ to cover T cell activation, co-inhibitory, T cell exhaustion, and T cell cytotoxicity gene signatures. The dot plots were created using ggplot2 R package, the heatmap was generated using ComplexHeatmap R package^52^. Expression of individual genes were plotted as bar graphs using GraphPad Prism.

### Tumor model and treatments

Procedures involving mice were performed in accordance with the standards of Fudan University and approved by Animal Ethical Committee of School of Pharmacy Fudan University (ethical permit 2023-08-SY-ZXY-82). Male NCG mice (20 ± 2 g body weight) were purchased from the GemPharmatech (Nanjing, China) and maintained under specific pathogen free conditions. NALM-6-luc cells (1 × 10^6^ cells) were intravenously injected into the mice at day 0. Tumor growth was monitored using bioluminescence imaging (BLI) system and tumor-bearing mice were randomized into the indicated groups (n = 5 per group) at day 3. Untransfected T cells, linear mRNA-CD19-CAR-T cells or cmRNA-CD19-CAR-T cells were intravenously injected into the corresponding mice (3 × 10^6^ cells per mouse) at days 3, 8, 13, and 18. Tumor burden was monitored twice a week by BLI system according to the instructions, and the total or mean photon number intensity of luciferase were quantified.

### Statistical analysis

Statistical analysis was performed by using GraphPad Prism software. Student’s t test and two-way ANOVA were used to determine the significance. Asterisks denote statistically significant p values (* p<0.05, ** p<0.01, *** p<0.001), and ns indicates no statistical significance (p > 0.05). Data were represented as the mean ± SD unless otherwise indicated.

## Results

### High-yield and scarless circRNA synthesis using an engineered PIE

Firstly, we engineered a new PIE platform based on Anabaena tRNALeu gene with high-yield and no extraneous nucleotide inserted into cmRNA product. The rationale behind the engineered PIE is to functionalize a selected 5’ end (termed as E2’) and 3’ end (termed as E1’) of the GOI as the functions of E2 and E1 of classical PIE, respectively. As illustrated in Fig. 1a, the scarless circRNA precursor in our engineered PIE constitutes of the 3’ half of ribozyme, the GOI (includes E2’ and E1’ surrogates at the ends of the GOI), and the 5’ half of ribozyme, this rational design enable us to synthesize really scarless circRNA products but keep the high yield advantages of classical PIE.

**Figure 1.**
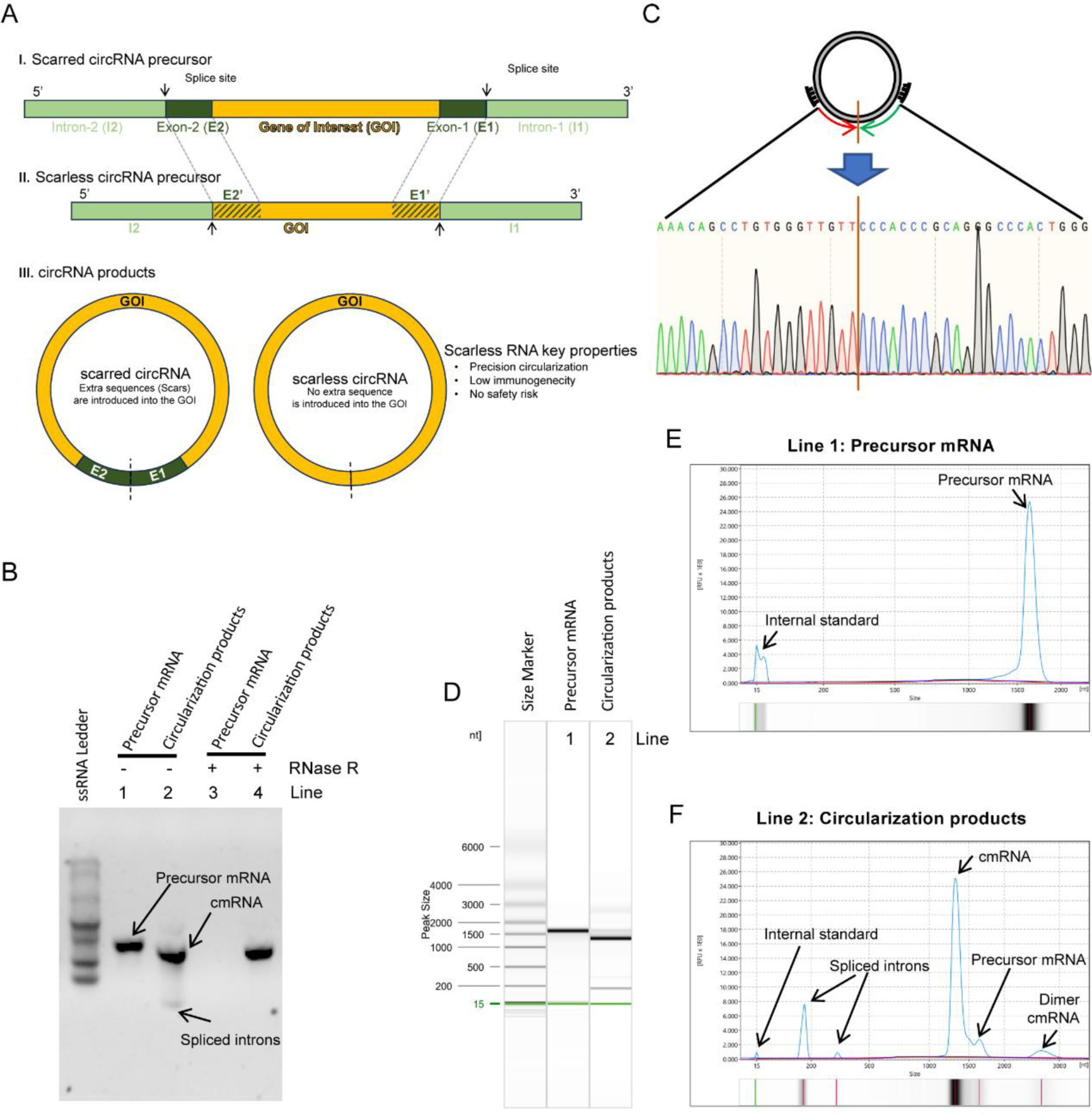
High-efficient synthesis of scarless circRNA using an engineered PIE-based circularization strategy. **A**, Engineering strategy for scarless circular RNA synthesis. Classical PIE strategy keeps E1 and E2 residues in circular RNA products (scar circRNA), while engineered new strategy generates scarless RNA products (scarless circRNA). **B**, Confirmation of the generation of circular format RNA molecules after circularization reactions using RNase R and RNase H treatments. **C**, Confirmation of precise sequence around the self-splicing junction site using sanger sequencing. **D**, **E** and **F**, Quantitative measurements of circularization reactions using capillary electrophoresis. The capillary electrophoresis separation results were shown as digit gel image bands (**D**) and classical peaks (**E** and **F**). **F**, The circularization efficiency was calculated by: 100% * the area of cmRNA peak / (the area of cmRNA peak + the area of precursor mRNA peak).

RNase R treatments for precursor mRNA and cmRNA product confirmed the circular format of products after circularization reactions which are resistant to RNase R while linear precursor mRNAs were digested (Fig. 1B, Fig. S1A-B). Circular mRNA products were also confirmed by Poly(A) polymerase (Fig. S1B) and RNase H endonuclease (Fig. S1C). Sanger sequencing indicated that the RNA sequence near the self-splicing junction site was identical to designed cmRNA (Fig. 1C and Fig. S1D), confirming the scarless capability of our engineered PIE. Capillary electrophoresis allows us to quantitatively measure the circularization yields and the results showed that the desired cmRNA accounted for ∼90% yield (Fig. S1E), which is comparable to classical PIE. The high yield of our engineered PIE is very friendly for downstream processes such as purification and scale-up. For example, the RNase R treatment was able to completely digest linear molecules and yield cmRNA with high purity (Fig. S1E). For simplicity, we named our engineered PIE as Hi-Scarless-PIE (high-yield and scarless PIE) thereafter.

### Robust protein expression of synthesized scarless cmRNAs *in vitro* and *in vivo*

Next, we examined the translation capability of scarless cmRNA produced by Hi-Scarless-PIE in cell line and in mouse. We synthesized cmRNAs using CVB3 IRES as translation initiation module to translate GLuc CDS (CVB3-GLuc for short). Linear GLuc mRNA (5’-capped, m1Ψ modified, and Ploy(A) tailed) and scarred CVB3-GLuc cmRNA (produced by PIE_176 nt scar version) were synthesized as controls (Fig. 2A). As Fig. 2B shown, both scarred and scarless cmRNAs exhibited prolonged protein expression durations over linear mRNA in HEK-293T cells. cmRNAs-based protein expression lasted for about 4.5 days to reach the so-called “half-life” (here means the roughly time point that the expression level was decreased to the half of the expression level at day 1), while the protein expression half-life of linear mRNA was about 2.5 days. The expression of GLuc was obviously detectable on day 7 for cmRNAs but was sharply diminished on day 3 and almost undetectable after day 4 for linear mRNAs. Obviously, the total amount of translated proteins over 7 days by cmRNAs were significantly more than that of linear mRNA (data not shown). These results demonstrated that cmRNA driven enhanced and prolonged protein expression *in vitro*. Moreover, we tested the protein expression capability of scarless cmRNA produced by Hi-Scarless-PIE for other GOIs including human erythropoietin (hEPO) and EGFP (Fig. 2C and D).

**Figure 2.**
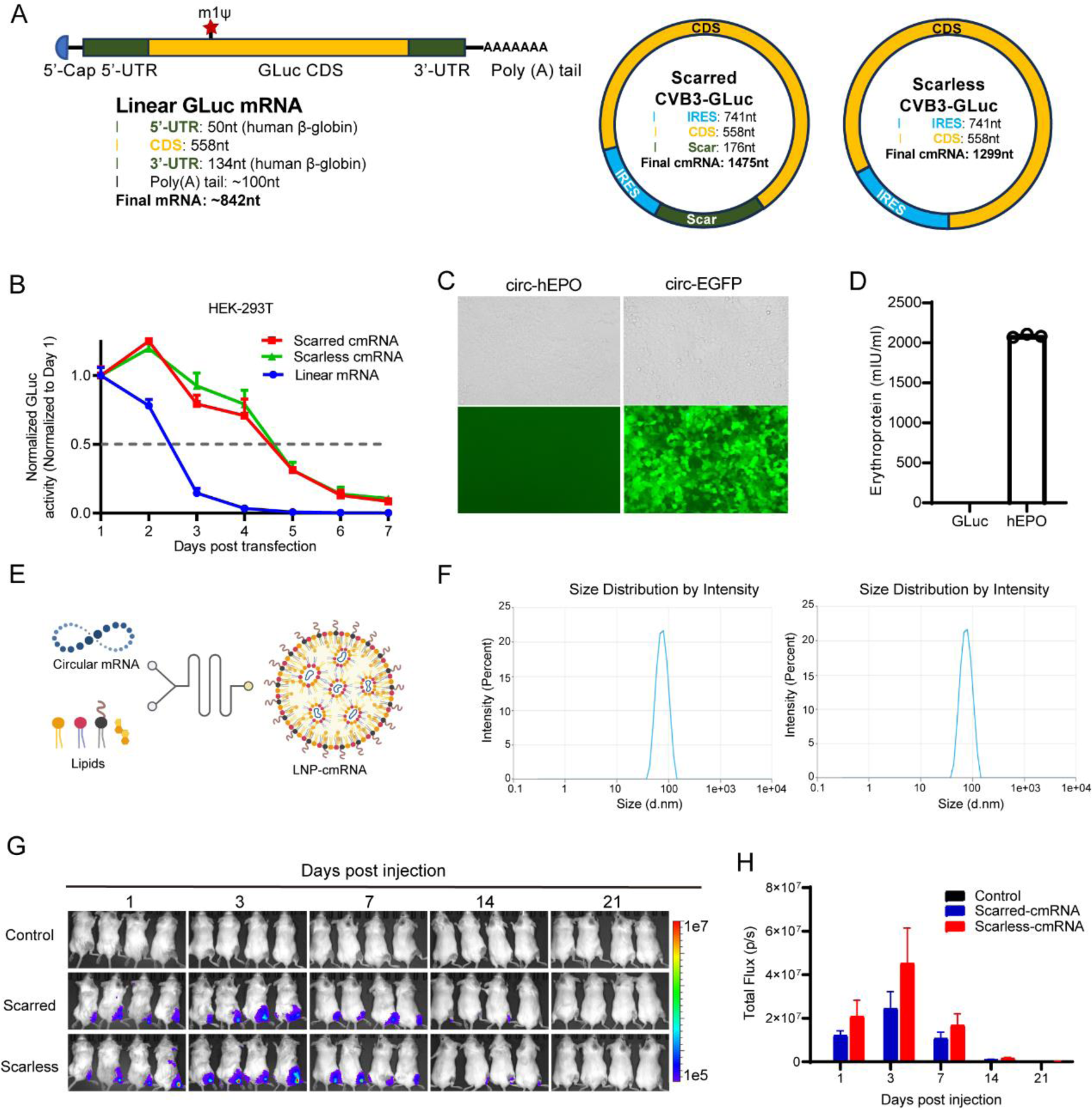
Robust protein expression of synthesized real-scarless cmRNAs *in vitro* and *in vivo*. **A,** Schematic illustrations depicting the architectural configurations of linear RNA, scarred circular RNA, and scarless circular RNA. **B**, Expression dynamics of GLuc mRNA in linear, scarred circular, and scarless circular mRNA formats in HEK-293T cells. Expressions of hEPO and EGFP scarless cmRNA in HEK-293T cells were observed under either fluorescent microscopy (**C**) or ELISA assays (**D**). **E**, The illustration of the formulation process of LNP encapsulating scarless cmRNA using microfluidics mixer. **F**, size distributions of LNP-cmRNA particles for scar cmRNA (left) and scarless cmRNA (right). **G**, Expressions of GLuc cmRNA in mice after intramuscular injection of 5µg of LNP-cmRNA per mouse were measured by bioluminescence imaging. **H**, The fluorescence intensity of mice in (**G**) were quantitatively presented. The error bars indicate SEM.

To test *in vivo* protein expression of scarless cmRNA, we formulated FLuc-coding cmRNAs with lipid nanoparticle (LNP) using microfluidics mixer (Fig. 2E). Formulated LNP-cmRNA exhibited appropriate average particle diameter size (∼75.0 nm), PDI (<0.15), Zeta-potential (−5 ∼5 mV), and encapsulation efficiency (∼90%) (Fig. 2f, Fig. S2). Intramuscular injection of 5µg of LNP-cmRNA resulted in potent FLuc protein expression for cmRNA and the expression lasted for 14 days (Fig. 2G). We observed that scarless cmRNA expressed slightly more FLuc proteins (on day 1, day 3, and day 7) compared with scarred cmRNA, even no statistical significance was detected due to large variations between individual mice (Fig. 2H).

### Scarless cmRNA exhibited enhanced and prolonged protein expression of CD19-CAR on primary human T cells

Using electroporation as the transfection method, we have prepared linear mRNA-based anti-CD19 CAR-T and cmRNA-based anti-CD19 CAR-T cells (Fig. 3A). CAR expression on the cell surface was monitored by incubating the cells with an PE-Labeled Human CD19 (20-291) Protein. Results indicated that CAR expression in the linear group peaked within 6 hours post-electroporation and began to decline after one day, reaching undetectable levels by the fourth day post-transfection. In contrast, CAR expression in the circular group reached its zenith a day after transfection and was sustained for two days, thereafter commencing a decline, with undetectable levels measured by the seventh day post-transfection. These data suggested that circular RNA confers a more sustained expression profile in T cells (Fig. 3B, C). Moreover, the median fluorescence intensity (MFI) of PE serves as an indicator of the intensity of CAR expression. The mean fluorescence intensity (MFI) of CAR in the linear cohort peaked 6 hours following transfection and subsequently exhibited a decline, reaching the levels of the untransfected cohort by the fourth day. In contrast, the circular cohort attained its acme 24 hours post transfection, surpassing the peak value of the linear cohort. This suggests a higher density of CAR on the T cells within the circular cohort compared to the linear one, thereby implying that the CAR-T cells of the circular group exhibited enhanced cytotoxic potential (Fig. 3D). In summary, the aforementioned results demonstrated that scarless cmRNA exhibited enhanced and prolonged protein expression of CD19-CAR in primary human T cells.

**Figure 3.**
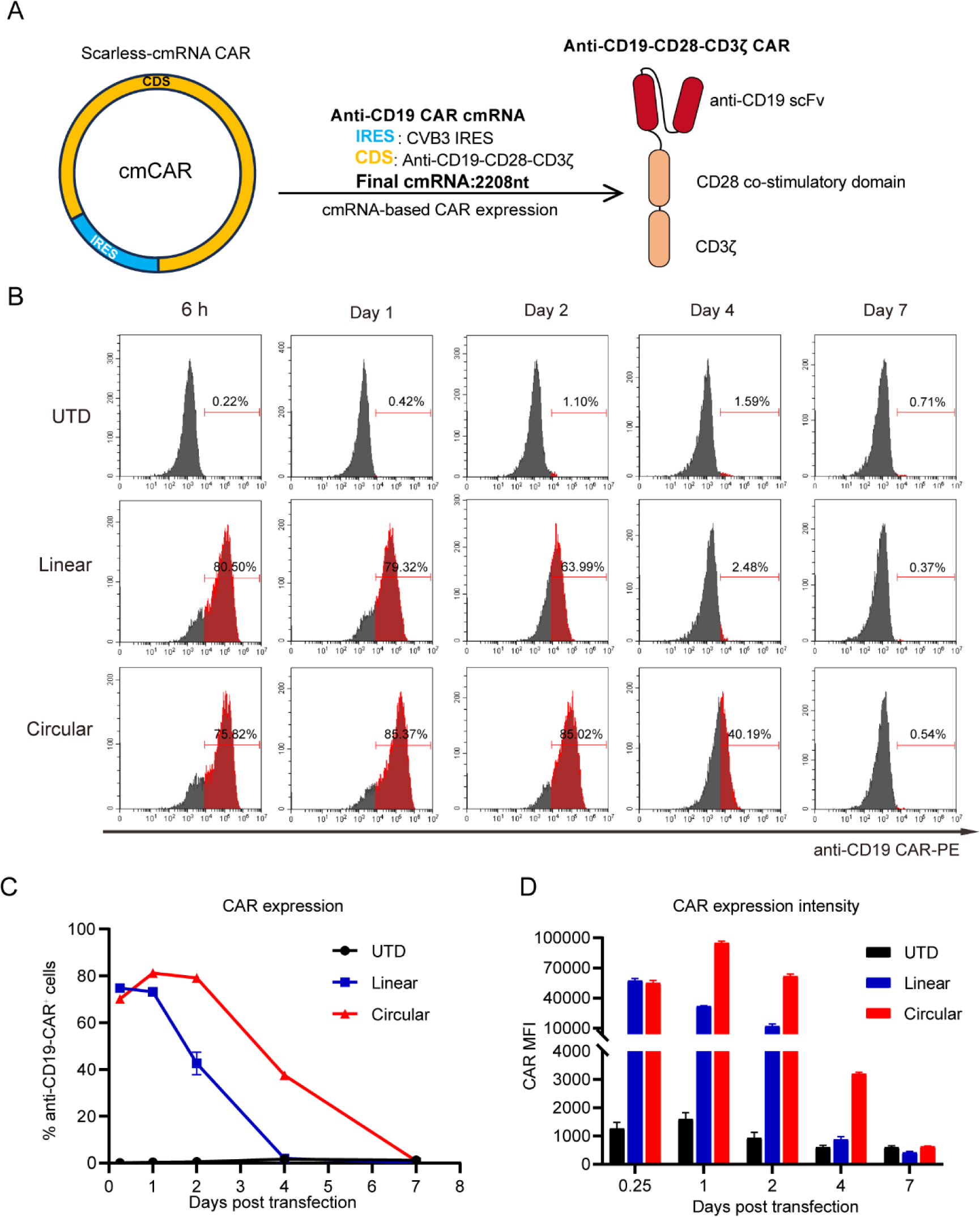
Scarless cmRNA exhibited enhanced and prolonged protein expression of CD19-CAR on primary human T cells. **A**, Structure of CAR-encoding cmRNA and the second-generation anti-CD19 CAR. **B**, Primary human T cells were activated, and transfected with linear mRNA (blue) or cmRNA (Red) at day 4. Representative histograms of CAR expression analysis from 6 hours to 7 days after transfection. PE-labeled human CD19 (20-291) Protein was used for CAR detection (CD19 CAR^+^) and Zombie Violet^TM^ Dye was performed to exclude dead cells. **C**, Proportion of CAR-expressing cells indicated by anti-CD19 CAR^+^ cells starting from 6 hours to 7 days post-transfection (n = 3 independent experiments/donors). **D**, The median intensity of CAR expression starting from 6 hours to 7 days post-transfection (n = 3 independent experiments).

### cmRNA-based anti-CD19 CAR-T cells shown potent cell-killing activities specifically against CD19^+^ malignant cells

To investigate the *in vitro* cytotoxicity of these two types of CAR-T cells (linear mRNA-based anti-CD19 CAR-T and cmRNA-based anti-CD19 CAR-T), we selected two CD19-positive cell lines NALM-6 and Raji, and one CD19-negative cell line K562 as target cells (Fig. S3). On the first day and fourth day following transfection, we co-cultured the CAR-T cells with the target cells at ratios of 1:1, 2:1, and 5:1 for 16 hours. Untransfected T cells were used as controls. The strengths and variations of the two types of CAR-T cells were assessed using flow cytometry. On the initial day following transfection, under diverse co-culture ratio conditions, both CAR-T cell variants were capable of nearly completely lysing CD19-positive neoplastic cells, with negligible cytotoxic effects observed against CD19-negative counterparts (Fig. 4A). Conversely, by the fourth day post-transfection, the linear mRNA-based anti-CD19 CAR-T cells displayed a cytolytic efficacy of 40%-60% against NALM-6 cells across varying co-culture ratios, while the circular mRNA-derived CAR-T cells achieved a cytotoxicity range of 60%-100%. In the context of Raji cells, the linear mRNA-based anti-CD19 CAR-T cells failed to demonstrate any cytotoxic impact under the tested conditions, whereas the cmRNA-based anti-CD19 CAR-T cells exhibited a lytic spectrum of 20%-50% (Fig. 4B). Collectively, these outcomes substantiated the superior cytotoxic potency of cmRNA-based anti-CD19 CAR-T cells against CD19-positive malignant cells, corroborating the findings from preceding experimental data.

**Figure 4.**
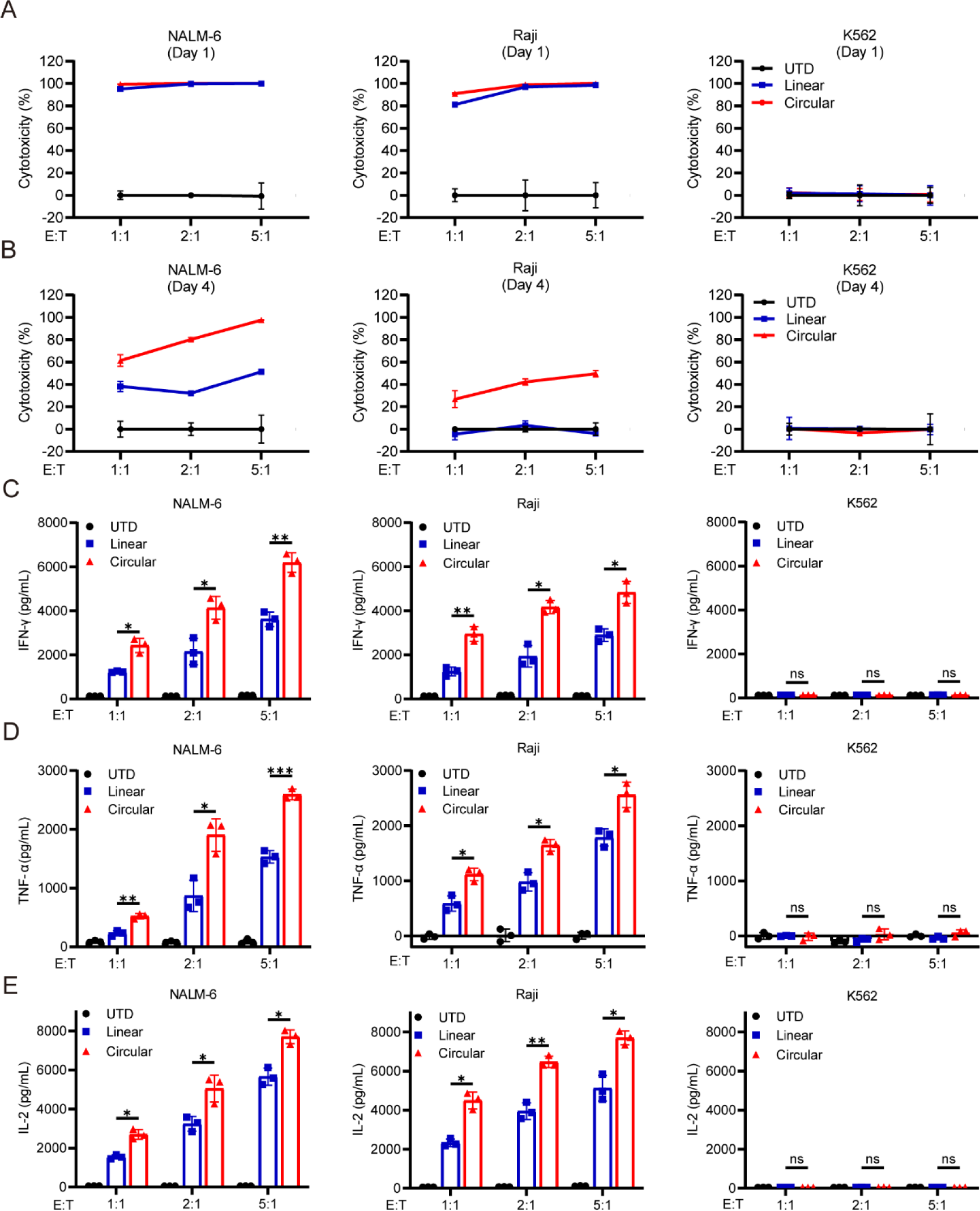
cmRNA-based anti-CD19 CAR-T cells shown potent target cell-killing activities with increased cytokines release. **A**, One day after transfection, linear mRNA-based anti-CD19 CAR-T or cmRNA-based anti-CD19 CAR-T cells were co-cultured with NALM-6 (left), Raji (middle) and K562 (right) cells at the E:T ratio of 1:1, 2:1 and 5:1 for 16h. **B**, Four days after transfection, linear mRNA-based anti-CD19 CAR-T or cmRNA-based anti-CD19 CAR-T were co-cultured with NALM-6, Raji and K562 cells at the E:T ratio of 1:1, 2:1 and 5:1 for. Untransfected (UTD) T cells were set as negative control. Bars represent mean ± SD of three independent experiments. **C**, IFN-γ release by linear mRNA-based anti-CD19 CAR-T or cmRNA-based anti-CD19 CAR-T cells after 16 h co-culture with NALM-6, Raji and K562 cells. **D**, TNF-ɑ release by linear mRNA-based anti-CD19 CAR-T or cmRNA-based anti-CD19 CAR-T cells after 16 h co-culture with NALM-6, Raji and K562 tumor cells. **E**, IL-2 release by linear mRNA-based anti-CD19 CAR-T or cmRNA-based anti-CD19 CAR-T cells after 16 h co-culture with NALM-6, Raji and K562 tumor cells. Data are presented as the mean ± SD (n=3 independent experiments; two-way ANOVA; *P < 0.05, **P < 0.01).

### Increased cytokines release of cmRNA-based CAR-T cells over mRNA-based counterparts

Next, we examined the level of interferon gamma (IFN-γ) and tumor necrosis factor alpha (TNF-α), key cytokines for effector cell function, and interleukin-2 (IL-2), a marker of CAR T-cell potency and persistence in CAR-T cells co-cultured with the indicated target cells. As illustrated in Figure 4C-E, untransfected T cells didn’t produce the aforementioned cytokines when co-cultured with tumor cells. Both linear mRNA-based anti-CD19 CAR-T and cmRNA-based anti-CD19 CAR-T cells, when co-cultured with CD19-negative tumor cells, also didn’t produce these cytokines. Only the co-culture of linear mRNA-based anti-CD19 CAR-T and cmRNA-based anti-CD19 CAR-T cells with CD19-positive tumor cells resulted in the dose-dependent release of substantial amounts of IFN-γ, TNF-α, and IL-2. Notably, the cmRNA-based anti-CD19 CAR-T cells produced significantly higher levels of all three cytokines compared to linear mRNA-based anti-CD19 CAR-T cells. This was consistent with the more potent cytotoxic capability of cmRNA-based anti-CD19 CAR-T cells against CD19-positive tumor cells.

### Dynamic changes of transcriptomics profiles of mCAR-T and cmCAR-T cells over time

Several studies have revealed that CAR-T cells produced by different ways^53^ and CAR-T cells in patients with difference outcomes (e.g., responders and long-remission patients) shown different transcriptomics profiles^50,54,55^. Therefore, we conducted RNA-seq experiments with the attempt to understand different cell-killing and cytokine release behaviors between mCAR-T and cmCAR-T cells from the transcription perspectives. Overall, untransfected T cells, cmCAR-T, and mCAR-T samples are obviously different with each other on day 2 and day 4, as indicated by the PCA dimension reduction analysis at transcriptomics level in Fig. 5A. The GZMB gene expression level of cmCAR-T cells was significantly higher than that of mCAR-T cells both on day 2 and day 4 (Fig.5B), which is in line with our previous results that cmCAR-T cells shown superior cytotoxic activities over mCAR-T cells (Fig. 4). The representative co-inhibitory genes PDCD1 and TIGIT shown different expression levels and changes (Fig. 5B). The absolute expression levels of PDCD1 (0.2-3.5 TPM) in all groups were much lower than that of TGIT (20-40 TPM). Transfection of mCAR and cmCAR resulted in increased expression of TIGIT on day 2 but decreased expression on day 4 compared with control baseline (Supplementary Table 2). This might suggest different transcriptomic profiles of CAR-T functional gene signatures between mCAR-T and cmCAR-T and on different time points.

**Figure 5.**
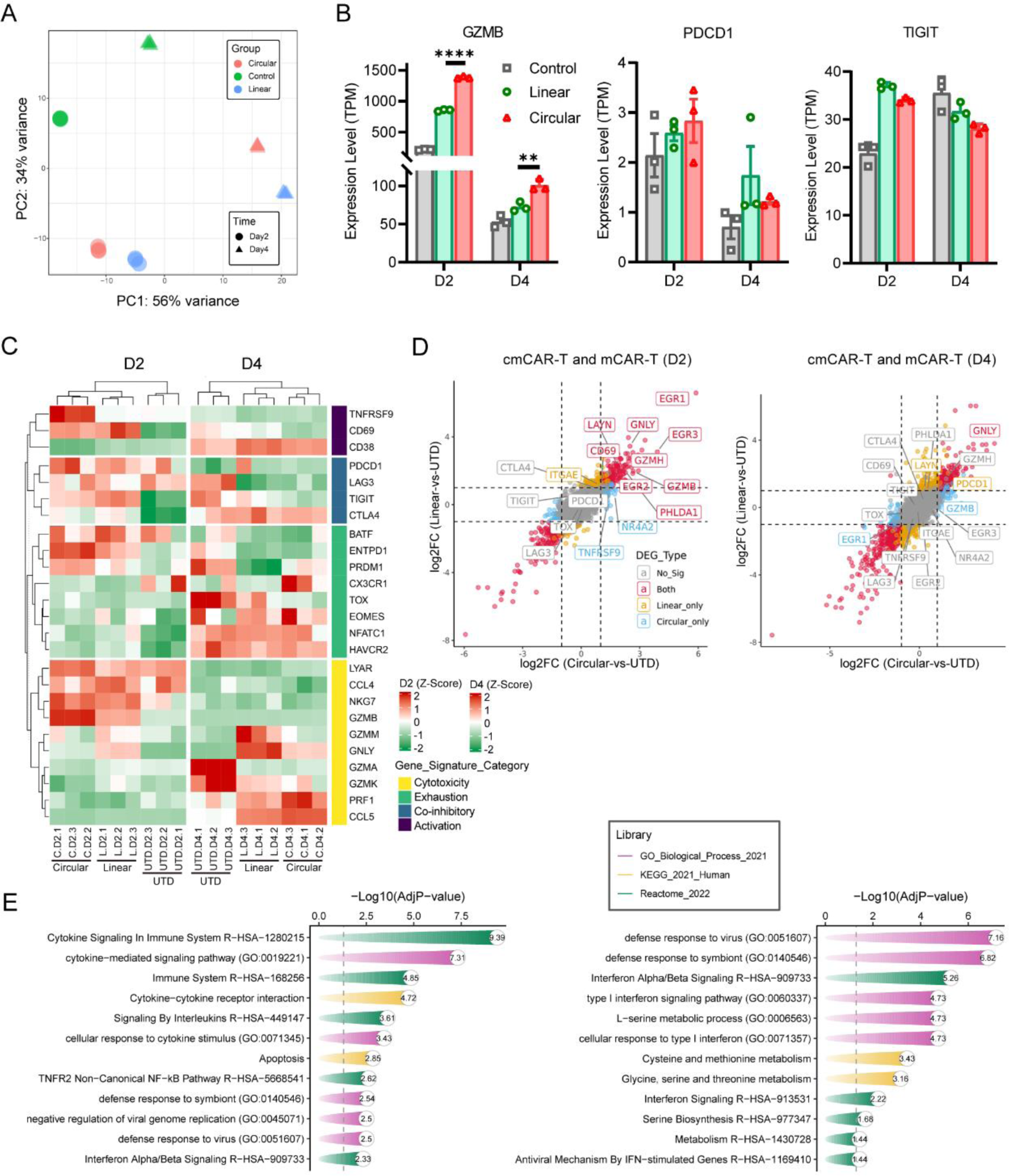
Transcriptomics profiles of mCAR-T and cmCAR-T cells on day 2 and day 4 post-transfection. **A**, Distribution of all samples under PCA dimensionality reduction analysis. **B**, The expression level (TPM) of GZMB and PDCD1 gene across groups. **C**, Expression spectrum of T cell activation, co-inhibitory, exhaustion, and cytotoxic gene signatures was visualized in heatmap. **D**, Differential expressed genes (DEGs) between cmCAR-T and UTD, and DEGs between mCAR-T and UTD were projected on x-axis and y-axis, respectively, according to their log2FC values. Representative genes involved in T cell activation, exhaustion, and cytotoxicity were labelled with gene names. **E**, Enriched biological processes and signaling pathways of upregulated DEGs in cmCAR-T compared to mCAR-T cells on day 2 and day 4 post-transfection. (**P < 0.01, ****P < 0.0001).

Therefore, we visualized well-documented gene signatures related to T cell activation, co-inhibitory signals, T-cell exhaustion, and T-cell cytotoxicity on heatmap (Fig. 5C). The dendrogram on the top of the heatmap shown that control T cell samples, mCAR-T samples, and cmCAR-T samples were clearly clustered as 3 clusters by heatmap hierarchical clustering algorithm, indicating obviously different overall patterns of these gene signatures between mCAR-T and cmCAR-T cells. To visualize the distribution of these gene signatures on the whole transcriptomics landscape, we performed DEG analysis and plotted all genes according to their log2FC in both mCAR-T-vs-Control and cmCAR-T-vs-Control DEG analysis. As shown in Fig. 5D, on day 2, there were 236 significantly upregulated DEGs in both mCAR-T and cmCAR-T, including cytotoxic genes (GZMB, GZMH) and T cell activation markers (CD69, GNLY, EGR1, EGR2, EGR3), while many T cell exhaustion and co-inhibitory signal genes (TIGIT, CTLA4, PDCD1, TOX, and LAG3) did not show significant changes in both mCAR-T and cmCAR-T, compared with control T cells, except for PHLDA1 and LAYN. However, many of those T cell activation and cytotoxicity related genes did not reach the DEG threshold on day 4, except for that GZMA was significantly upregulated only in cmCAR-T cells, and LAYN (a T-cell exhaustion marker) was significantly upregulated only in mCAR-T cells. To further understand the transcription differences between mCAR-T and cmCAR-T cells, we directly compared cmCAR-T cells with mCAR-T cells using DEG analysis. There were 113 and 232 upregulated DEGs in cmCAR-T cells on day 2 and day 4, respectively. Enrichment analysis indicated that these DEGs were significantly overrepresented in several cytokine signaling pathways on day 2 and interferon signaling on day 4 in cmCAR-T cells (Fig. 5E), which in agreement with previous data (Fig. 4C-E, and Supplementary Table 2).

Collectively, these transcription level evidence suggested that T cell activation and cytotoxic signals might be different between mCAR-T and cmCAR-T cells, and CAR-T functional signals were transient in both mCAR-T and cmCAR-T cells, but cmCAR-T cells exhibited enhanced and prolonged cytotoxicity over mCAR-T.

### cmRNA-based anti-CD19 CAR-T cells eliminated NALM-6 cells in mouse model

To further investigate the *in vivo* persistence of the two types of CAR-T cells, we administered the CAR-T cells one day post-electroporation via tail vein injection at a dosage of 3×10^6^ cells per mouse (Fig. 6A). Blood samples were collected at day 1, day 3, and day 5 post-injection to assess the duration of CAR-T cell presence within the mice. As depicted in Figure 6B, CAR-T cells were detectable on the first day post-injection, with the linear mRNA group exhibiting a proportion of 40%, while the circular mRNA group surpassed 80%. By the third day post-injection, the proportion of linear mRNA CAR-T cells had dwindled to 20%, whereas the circular mRNA group maintained over 50%. These findings are in concordance with the previous *in vitro* results, indicating that cmRNA-based anti-CD19 CAR-T cells exhibited a more prolonged persistence *in vivo*. Subsequently, we evaluated the therapeutic timing and dosage for the cmRNA group. As illustrated in Figure 6C, 1×10^6^ luciferase-labeled NALM-6 (NALM-6-luc) cells were injected into each mouse on day 0. Tumor proliferation within the mice was monitored through bioluminescence imaging, with the imaging conducted on the third day, marking the commencement of treatment. The low-dose and high-dose group received an injection of 1×10^6^ and 3×10^6^ cmRNA-based anti-CD19 CAR-T cells per mouse, respectively. The treatment was conducted every four days for three times. Bioluminescence imaging outcomes indicated that the tumor burden in both the low-dose and high-dose treatment cohorts was significantly reduced compared to the untransfected group. Following three rounds of treatment, a trend of increasing tumor proliferation was observed within the low-dose group, with one particular instance of growth noted. Moreover, the high-dose treatment demonstrated a more efficacious tumor clearance compared to the low-dose regimen (Fig. 6D-F). These findings suggested that the dosage regimen of 3×10^6^ CAR-T cells coupled with an increased frequency of treatments may be warranted.

**Figure 6.**
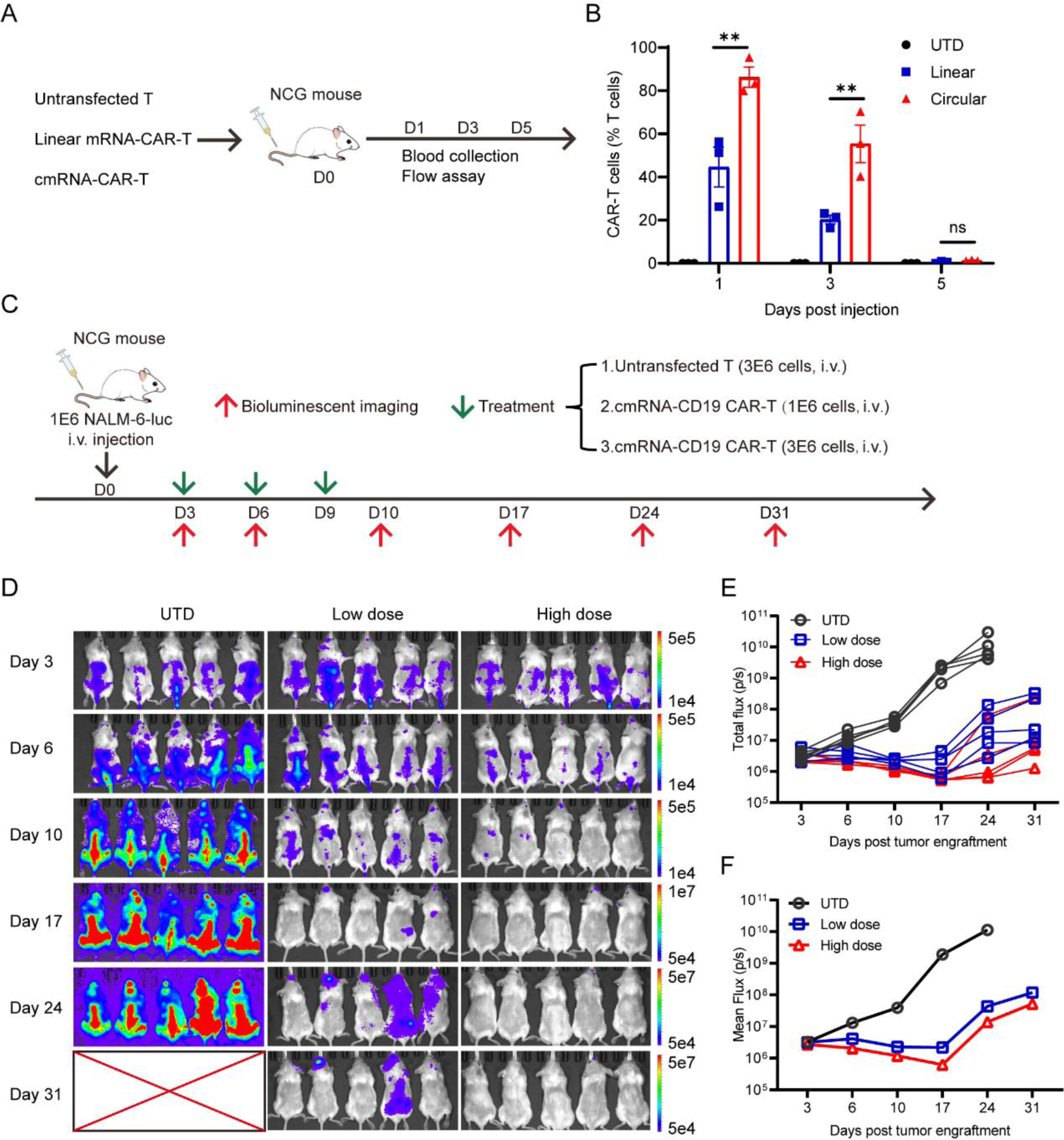
cmRNA-based anti-CD19 CAR-T cells eliminated NALM-6 cells *in vivo*. (**A**, **B**) Schematic diagram and detection of the persistence of CAR-T cells within mice using flow cytometry. **C**, Schematic diagram of the NALM-6 tumor model and procedure of linear mRNA-based anti-CD19 CAR-T or cmRNA-based anti-CD19 CAR-T cells infusion time and dose. **D**, Bioluminescence imaging analysis of NALM-6 tumors, n = 5 mice per group. (**E**, **F**) The tumor burden was quantified as the total flux (each line represents a single mouse) or mean flux from luciferase intensity of each mouse. (ns, not significant; **P < 0.01).

### cmRNA-based anti-CD19 CAR-T cells elicited potent antitumor efficacy *in vivo*

We next assessed the feasibility and efficacy of linear mRNA-based anti-CD19 CAR-T cells and cmRNA-based anti-CD19 CAR-T cells in NALM-6 tumor model. As illustrated in Figure 7A, 1 × 10^6^ NALM-6-luc cells were transplanted into NCG mouse on day 0, and 3 ×10^6^ untransfected T cells or linear mRNA-based anti-CD19 CAR-T cells or cmRNA-based anti-CD19 CAR-T cells were injected on day 3, bioluminescence imaging was performed for detection and quantification after tumor cell inoculation. The bioluminescence imaging analysis revealed that the tumor burden of mice infused with cmRNA-based anti-CD19 CAR-T cells was significantly lower than those of the untransfected T cells and linear mRNA-based anti-CD19 CAR-T cells (Fig. 7B-D). Moreover, significant body weight loss was observed in the control groups due to the severe tumor burden, whereas a relatively stable body weight was observed in the groups infused with cmRNA-based anti-CD19 CAR-T cells (Fig. S4). Besides, infusions cmRNA-based anti-CD19 CAR-T cells significantly prolonged the survival time of the mice (Fig. 7E). On day 12, blood samples were collected to detect CAR-T cells and data showed that the proportion of CAR-T cells in the circular group was significantly higher than that in the linear group (Fig. 7F). On day 20 of the experiment, analysis of memory T cells showed that the proportion of memory T cells in the circular group was significantly higher than that in the linear group, indicating the persistent antitumor efficacy of cmRNA-based anti-CD19 CAR-T cells (Fig. 7G). Collectively, the above results proved that cmRNA-based anti-CD19 CAR-T cells could efficiently eliminate NALM-6 cells *in vivo* and provide long-lasting antitumor efficacy.

**Figure 7.**
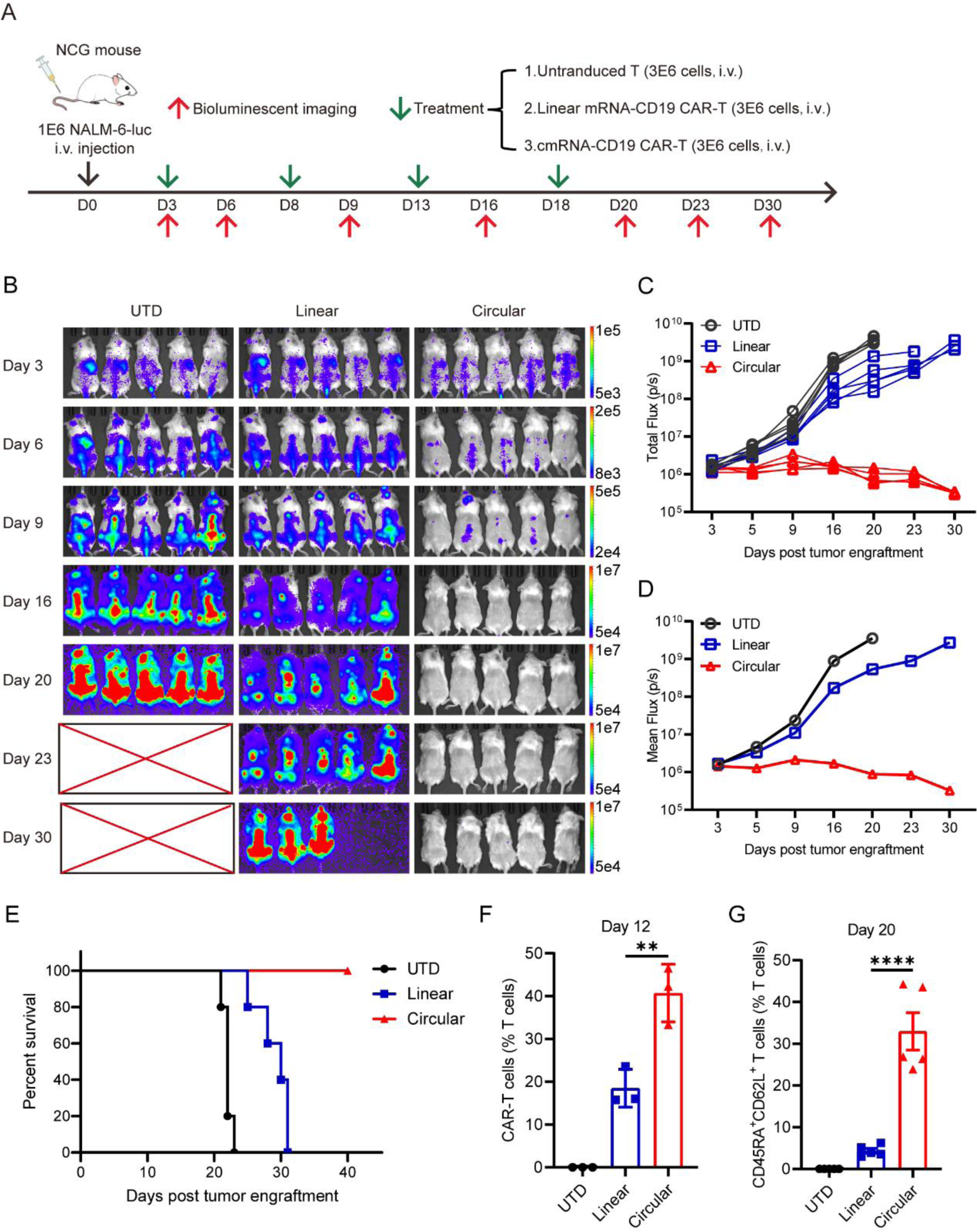
cmRNA-based CAR-T cells outperformed mRNA-based CAR-T cells in eliminating target cells *in vivo*. **A**, Schematic diagram of NALM-6 tumor model and procedure of linear mRNA-based anti-CD19 CAR-T or cmRNA-based anti-CD19 CAR-T cells infusion time and dose. (**B**-**D**) Bioluminescence imaging analysis of NALM-6 tumor, n = 5 mice per group. Tumor burden was quantified as the total flux (**C**, each line represents single mouse) or mean flux (**D**) from luciferase intensity of each mouse. **E**, Survival curve analysis of mice in the different treatment groups. **F**, On the twelfth day of the experiment, the proportion of anti-CD19 CAR-T cells was determined by flow cytometry. **G**, On the twenty-second day of the experiment, the proportion of memory T cells was assessed by flow cytometry. (**P < 0.01, ***P < 0.001).

## Discussion

Several CAR-T cell therapies have been approved for hematological malignancies and ongoing investigations are expanding CAR-T to solid tumors and other indications beyond cancer^1,2^. However, the extensive clinical applications have raised a series of safety concerns including the risk for secondary T-cell malignancies following CD19-directed or BCMA-directed CAR-T cell therapies, which may result from the inaccuracy transgene integration of CAR. The FDA explicitly stated that, with all gene therapy products using integrating vectors, the potential risk of developing secondary malignancies is categorized as a class warning^56^. Besides the dangerous safety risk, viral vector-based manufacturing of CAR-T has a relatively high incidence of side effects such as CRS. A safer version of CAR-T is urgently needed to minimize safety issues and more importantly, unleash the full potentials of CAR-T cell therapy in noncancerous indications which is extremely sensitive to risk-benefit balance^1,11,17^.

mRNA platform offers an ideal strategy to avoid the risks stem from viral vectors, genome integration, and permanent transgene expression of traditional CAR-T technologies^21^. Indeed, *ex vivo* mRNA-based CAR-T cells have been tested for an extended period in early-stage of clinical trials. Along with the advances in targeted LNP delivery systems, the *in vivo* CAR-T therapy, which generates CAR-T cells in human body via targeted delivering CAR-encoding mRNA into T cells, will be a gamechanger in the wave of developing next generation of CAR-T therapies^23,24^. However, the intrinsically unstable nature of linear mRNA compromised the efficacy of mRNA-based CAR-T due to short-term expression duration.

Since the first report of high-efficient *in vitro* synthesis of cmRNA in 2018^35^, circular RNA gains a lot of attentions as an updated mRNA platform for therapeutic purposes^37,40,57^. In this study, we developed a new PIE strategy to synthesize cmRNA with scarless and high-yield capabilities, which are very important druggability considerations for drug development. We confirmed that cmRNA increases the amount and extends the duration of CAR expression on human T cells. Subsequently, we demonstrated *in vitro* and *in vivo* advantages of cmRNA in CAR expression and CAR-T therapy using an approved anti-CD19-CAR construct as an example. cmRNA-based anti-CD19 CAR-T cells showed increased intensity and prolonged duration of CAR expression on primary human T cells. Indeed, our results demonstrated that primary human T cells transfected with CAR-encoding cmRNA were more effective at priming the cytotoxic T cells compared to those transfected with linear mRNA *in vitro*. Compared with linear mRNA, the intrinsic stability and long expression half-life of cmRNA represent additional advantages in mRNA-based CAR-T therapies, given the fact that CAR-encoding mRNA molecules will be divided into daughter CAR-T cells along with the CAR-T cell expansion when interacting with targeted cells *in vivo*.

Collectively, we have engineered a PIE platform to synthesize high-yield and scarless cmRNA for potent CAR expression and long-lasting CAR-T efficacy. Compared to linear mRNA counterparts, cmRNA-based anti-CD19 CAR-T cells elicited superior anti-tumor efficacy, demonstrated by parallel lines of evidence including *in vitro* specific cell-killing, cytokine release, transcriptomics patterns, and *in vivo* tumor elimination and survival benefits. Our findings indicated that cmRNA-based CAR-T could efficiently eliminate target cells and provide long-lasting antitumor efficacy, providing a potent platform for mRNA-based CAR-T cell therapies. Together with future advances in T cell-targeted delivery systems, cmRNA will further unleashing full potentials of mRNA technologies in CAR-based *in vivo* cell therapies.

## Reference

1 Baker, D. J., Arany, Z., Baur, J. A., Epstein, J. A. & June, C. H. CAR T therapy beyond cancer: the evolution of a living drug. Nature 619, 707–715, doi:10.1038/s41586-023-06243-w (2023).

2 Labanieh, L. & Mackall, C. L. CAR immune cells: design principles, resistance and the next generation. Nature 614, 635–648, doi:10.1038/s41586-023-05707-3 (2023).

3 Mackensen, A. et al. Anti-CD19 CAR T cell therapy for refractory systemic lupus erythematosus. Nat Med 28, 2124–2132, doi:10.1038/s41591-022-02017-5 (2022).

4 Wang, X. et al. Allogeneic CD19-targeted CAR-T therapy in patients with severe myositis and systemic sclerosis. Cell, doi:10.1016/j.cell.2024.06.027 (2024).

5 Krickau, T. et al. CAR T-cell therapy rescues adolescent with rapidly progressive lupus nephritis from haemodialysis. Lancet 403, 1627–1630, doi:10.1016/s0140-6736(24)00424-0 (2024).

6 Rurik, J. G. et al. CAR T cells produced in vivo to treat cardiac injury. Science 375, 91–96, doi:10.1126/science.abm0594 (2022).

7 Amor, C. et al. Senolytic CAR T cells reverse senescence-associated pathologies. Nature 583, 127–132, doi:10.1038/s41586-020-2403-9 (2020).

8 Yang, D. et al. NKG2D-CAR T cells eliminate senescent cells in aged mice and nonhuman primates. Sci Transl Med 15, eadd1951, doi:10.1126/scitranslmed.add1951 (2023).

9 Amor, C. et al. Prophylactic and long-lasting efficacy of senolytic CAR T cells against age-related metabolic dysfunction. Nat Aging 4, 336–349, doi:10.1038/s43587-023-00560-5 (2024).

10 Morris, E. C., Neelapu, S. S., Giavridis, T. & Sadelain, M. Cytokine release syndrome and associated neurotoxicity in cancer immunotherapy. Nat Rev Immunol 22, 85–96, doi:10.1038/s41577-021-00547-6 (2022).

11 Brudno, J. N. & Kochenderfer, J. N. Current understanding and management of CAR T cell-associated toxicities. Nat Rev Clin Oncol 21, 501–521, doi:10.1038/s41571-024-00903-0 (2024).

12 Suran, M. FDA Adds Boxed Warning to CAR T-Cell Therapies, but Says Benefits Outweigh Risks of Secondary Cancers. Jama 331, 818–820, doi:10.1001/jama.2024.1011 (2024).

13 Ghilardi, G. et al. T cell lymphoma and secondary primary malignancy risk after commercial CAR T cell therapy. Nat Med 30, 984–989, doi:10.1038/s41591-024-02826-w (2024).

14 Verdun, N. & Marks, P. Secondary Cancers after Chimeric Antigen Receptor T-Cell Therapy. N Engl J Med 390, 584–586, doi:10.1056/NEJMp2400209 (2024).

15 Levine, B. L. et al. Unanswered questions following reports of secondary malignancies after CAR-T cell therapy. Nat Med 30, 338–341, doi:10.1038/s41591-023-02767-w (2024).

16 Elsallab, M. et al. Second primary malignancies after commercial CAR T-cell therapy: analysis of the FDA Adverse Events Reporting System. Blood 143, 2099–2105, doi:10.1182/blood.2024024166 (2024).

17 Rafiq, S., Hackett, C. S. & Brentjens, R. J. Engineering strategies to overcome the current roadblocks in CAR T cell therapy. Nat Rev Clin Oncol 17, 147–167, doi:10.1038/s41571-019-0297-y (2020).

18 Zhang, J. et al. Non-viral, specifically targeted CAR-T cells achieve high safety and efficacy in B-NHL. Nature 609, 369–374, doi:10.1038/s41586-022-05140-y (2022).

19 Foy, S. P. et al. Non-viral precision T cell receptor replacement for personalized cell therapy. Nature 615, 687–696, doi:10.1038/s41586-022-05531-1 (2023).

20 Parayath, N. N., Stephan, S. B., Koehne, A. L., Nelson, P. S. & Stephan, M. T. In vitro-transcribed antigen receptor mRNA nanocarriers for transient expression in circulating T cells in vivo. Nat Commun 11, 6080, doi:10.1038/s41467-020-19486-2 (2020).

21 Wu, J., Wu, W., Zhou, B. & Li, B. Chimeric antigen receptor therapy meets mRNA technology. Trends Biotechnol 42, 228–240, doi:10.1016/j.tibtech.2023.08.005 (2024).

22 Liu, C. et al. mRNA-based cancer therapeutics. Nat Rev Cancer 23, 526–543, doi:10.1038/s41568-023-00586-2 (2023).

23 Parhiz, H., Atochina-Vasserman, E. N. & Weissman, D. mRNA-based therapeutics: looking beyond COVID-19 vaccines. Lancet 403, 1192–1204, doi:10.1016/s0140-6736(23)02444-3 (2024).

24 Short, L., Holt, R. A., Cullis, P. R. & Evgin, L. Direct in vivo CAR T cell engineering. Trends Pharmacol Sci 45, 406–418, doi:10.1016/j.tips.2024.03.004 (2024).

25 Chung, J. B., Brudno, J. N., Borie, D. & Kochenderfer, J. N. Chimeric antigen receptor T cell therapy for autoimmune disease. Nat Rev Immunol, doi:10.1038/s41577-024-01035-3 (2024).

26 Barrett, D. M. et al. Treatment of advanced leukemia in mice with mRNA engineered T cells. Hum Gene Ther 22, 1575–1586, doi:10.1089/hum.2011.070 (2011).

27 Tombácz, I. et al. Highly efficient CD4+ T cell targeting and genetic recombination using engineered CD4+ cell-homing mRNA-LNPs. Mol Ther 29, 3293–3304, doi:10.1016/j.ymthe.2021.06.004 (2021).

28 Foster, J. B. et al. Development of GPC2-directed chimeric antigen receptors using mRNA for pediatric brain tumors. J Immunother Cancer 10, doi:10.1136/jitc-2021-004450 (2022).

29 Tilsed, C. M. et al. IL7 increases targeted lipid nanoparticle-mediated mRNA expression in T cells in vitro and in vivo by enhancing T cell protein translation. Proc Natl Acad Sci U S A 121, e2319856121, doi:10.1073/pnas.2319856121 (2024).

30 Beatty, G. L. et al. Mesothelin-specific chimeric antigen receptor mRNA-engineered T cells induce anti-tumor activity in solid malignancies. Cancer Immunol Res 2, 112–120, doi:10.1158/2326-6066.Cir-13-0170 (2014).

31 Beatty, G. L. et al. Activity of Mesothelin-Specific Chimeric Antigen Receptor T Cells Against Pancreatic Carcinoma Metastases in a Phase 1 Trial. Gastroenterology 155, 29–32, doi:10.1053/j.gastro.2018.03.029 (2018).

32 Shah, P. D. et al. Phase I Trial of Autologous RNA-electroporated cMET-directed CAR T Cells Administered Intravenously in Patients with Melanoma and Breast Carcinoma. Cancer Res Commun 3, 821–829, doi:10.1158/2767-9764.Crc-22-0486 (2023).

33 Liu, C. X. & Chen, L. L. Circular RNAs: Characterization, cellular roles, and applications. Cell 185, 2016–2034, doi:10.1016/j.cell.2022.04.021 (2022).

34 Metkar, M., Pepin, C. S. & Moore, M. J. Tailor made: the art of therapeutic mRNA design. Nat Rev Drug Discov 23, 67–83, doi:10.1038/s41573-023-00827-x (2024).

35 Wesselhoeft, R. A., Kowalski, P. S. & Anderson, D. G. Engineering circular RNA for potent and stable translation in eukaryotic cells. Nat Commun 9, 2629, doi:10.1038/s41467-018-05096-6 (2018).

36 Wesselhoeft, R. A. et al. RNA Circularization Diminishes Immunogenicity and Can Extend Translation Duration In Vivo. Mol Cell 74, 508–520.e504, doi:10.1016/j.molcel.2019.02.015 (2019).

37 Chen, R. et al. Engineering circular RNA for enhanced protein production. Nat Biotechnol 41, 262–272, doi:10.1038/s41587-022-01393-0 (2023).

38 Choi, S. W. & Nam, J. W. Optimal design of synthetic circular RNAs. Exp Mol Med 56, 1281–1292, doi:10.1038/s12276-024-01251-w (2024).

39 Gholamalipour, Y., Karunanayake Mudiyanselage, A. & Martin, C. T. 3’ end additions by T7 RNA polymerase are RNA self-templated, distributive and diverse in character-RNA-Seq analyses. Nucleic Acids Res 46, 9253–9263, doi:10.1093/nar/gky796 (2018).

40 Guo, S. K. et al. Therapeutic application of circular RNA aptamers in a mouse model of psoriasis. Nat Biotechnol, doi:10.1038/s41587-024-02204-4 (2024).

41 Liu, C. X. et al. RNA circles with minimized immunogenicity as potent PKR inhibitors. Mol Cell 82, 420–434.e426, doi:10.1016/j.molcel.2021.11.019 (2022).

42 Chen, Y. G. et al. N6-Methyladenosine Modification Controls Circular RNA Immunity. Mol Cell 76, 96–109.e109, doi:10.1016/j.molcel.2019.07.016 (2019).

43 Eisenstein, M., Garber, K., Landhuis, E. & DeFrancesco, L. Nature Biotechnology’s academic spinouts 2021. Nat Biotechnol 40, 1551–1562, doi:10.1038/s41587-022-01530-9 (2022).

44 Lee, K. H. et al. Efficient circular RNA engineering by end-to-end self-targeting and splicing reaction using Tetrahymena group I intron ribozyme. Mol Ther Nucleic Acids 33, 587–598, doi:10.1016/j.omtn.2023.07.034 (2023).

45 Cui, J. et al. A precise and efficient circular RNA synthesis system based on a ribozyme derived from Tetrahymena thermophila. Nucleic Acids Res 51, e78, doi:10.1093/nar/gkad554 (2023).

46 Dobin, A. et al. STAR: ultrafast universal RNA-seq aligner. Bioinformatics 29, 15–21, doi:10.1093/bioinformatics/bts635 (2013).

47 Liao, Y., Smyth, G. K. & Shi, W. featureCounts: an efficient general purpose program for assigning sequence reads to genomic features. Bioinformatics 30, 923–930, doi:10.1093/bioinformatics/btt656 (2014).

48 Love, M. I., Huber, W. & Anders, S. Moderated estimation of fold change and dispersion for RNA-seq data with DESeq2. Genome Biol 15, 550, doi:10.1186/s13059-014-0550-8 (2014).

49 Evangelista, J. E. et al. Enrichr-KG: bridging enrichment analysis across multiple libraries. Nucleic Acids Res 51, W168–w179, doi:10.1093/nar/gkad393 (2023).

50 Anderson, N. D. et al. Transcriptional signatures associated with persisting CD19 CAR-T cells in children with leukemia. Nat Med 29, 1700–1709, doi:10.1038/s41591-023-02415-3 (2023).

51 Louie, R. H. Y. et al. CAR(+) and CAR(-) T cells share a differentiation trajectory into an NK-like subset after CD19 CAR T cell infusion in patients with B cell malignancies. Nat Commun 14, 7767, doi:10.1038/s41467-023-43656-7 (2023).

52 Gu, Z. Complex heatmap visualization. Imeta 1, e43, doi:10.1002/imt2.43 (2022).

53 Niu, A. et al. Differences in the phenotypes and transcriptomic signatures of chimeric antigen receptor T lymphocytes manufactured via electroporation or lentiviral transfection. Front Immunol 14, 1068625, doi:10.3389/fimmu.2023.1068625 (2023).

54 Haradhvala, N. J. et al. Distinct cellular dynamics associated with response to CAR-T therapy for refractory B cell lymphoma. Nat Med 28, 1848–1859, doi:10.1038/s41591-022-01959-0 (2022).

55 Rezvan, A. et al. Identification of a clinically efficacious CAR T cell subset in diffuse large B cell lymphoma by dynamic multidimensional single-cell profiling. Nat Cancer 5, 1010–1023, doi:10.1038/s43018-024-00768-3 (2024).

56 Zhu, Z. et al. mRNA-Engineered CD5-CAR-γδTCD5-Cells for the Immunotherapy of T-Cell Acute Lymphoblastic Leukemia. Adv Sci, e2400024. doi: 10.1002/advs.202400024. (2024).

57 Qu, L. et al. Circular RNA vaccines against SARS-CoV-2 and emerging variants. Cell 185, 1728–1744.e1716, doi:10.1016/j.cell.2022.03.044 (2022).

